# Frequency selective encoding of substrate vibrations in the somatosensory cortex

**DOI:** 10.1101/264747

**Authors:** Mario Prsa, Daniel Huber

## Abstract

Sensing vibrations that propagate through solid substrates conveys fundamental information about moving objects and other nearby dynamic events. Here we report that neurons responsive to substrate vibrations applied to the mouse forelimb reveal a new way of representing frequency information in the primary somatosensory cortex (S1). In contrast to vibrotactile stimulation of primate glabrous skin, which produces temporally entrained spiking and frequency independent firing rates, we found that mouse S1 neurons rely on a different coding scheme: their spike rates are conspicuously tuned to a preferred frequency of the stimulus. Histology, peripheral nerve block and optogenetic tagging experiments furthermore reveal that these responses are associated with the activation of mechanoreceptors located in deep subdermal tissue of the distal forelimb. We conclude that the encoding of frequency information of substrate-borne vibrations in the mouse S1 might be analogous to the representation of pitch of airborne sound in auditory cortex.

## Introduction

Vibrations are ecologically relevant sensory stimuli that can appear in different forms and their spectral content can convey different types of information. For example, the frequency of skin vibrations produced by actively displacing a fingertip across an object provides information about the surface texture. Spikes of S1 neurons responding to skin vibrations of primate fingertips are found to be temporally locked to a preferred phase of the oscillatory cycle, but their rates are reported to be insensitive to stimulus frequency (Harvey et al., 2013; Mountcastle et al., 1969). Frequency dependent spike rate modulations could only be obtained with pulsed vibrations in the flutter range (<40 Hz) (Salinas et al., 2000), in which individual stimulus cycles and not pitch dominate the percept (Saal et al., 2016). The same seems to be true in rodents, where active whisker-mediated sensation has been advanced as the main modus operandi for collecting somatosensory information from the environment (Diamond et al., 2008); a functional equivalent to active hand palpations in primates. Phase-locked spiking and frequency independent firing rates of neural responses to passive sinusoidal vibrations of single whiskers have been reported in barrel cortex (Ewert et al., 2008) (but see (Arabzadeh et al., 2004; Arabzadeh et al., 2003)). It therefore seems that in such active sensory systems frequencies are not discriminated based on differences in rates of discharge of cortical neurons. Instead, the proposed neural coding scheme for frequency recognition is based on identifying the dominant phase of the oscillatory stimulus in the cyclically entrained spike trains.

Different types of vibratory stimuli are generated by external movements and they can be perceived passively. These are transmitted through the solid substrate, and their frequency might indicate the size, speed or distance of, for instance, a nearby conspecific, predator or prey. This passive sensory channel is therefore not primarily adapted to identify object textures or shapes, but rather tuned for gathering essential information about events occurring in the nearby environment. Substrate vibrations propagate away from the surface of contact, typically through the limbs (Hunt, 1961; Hunt and McIntyre, 1960; O’Connell-Rodwell, 2007) and might therefore activate a broad set of mechanoreceptors distributed along its bones, joints and possibly even in more distant body parts (Bell et al., 1994; Hunt, 1961; Zelena, 1994). The involved network of mechanoreceptors might thus be very different from the one probed in experiments delivering local skin vibrations at the fingertip, since such vibrations attenuate exponentially with distance and do not seem to propagate beyond the finger (Manfredi et al., 2012). If substrate vibrations indeed rely on a different functional network of mechanoreceptors, might they also have a distinct neural representation of vibration frequency in the cortex?

To address this question we sought to characterize how neurons in the mouse forelimb representation of the somatosensory cortex (fS1) encode the frequency of substrate vibrations. Using two-photon calcium imaging and electrophysiology, we found that neurons in the mouse fS1 show spike rates tuned to different stimulus frequencies and lack temporally entrained spiking. They bandpass filter the mechanical stimuli to favor a narrow bandwidth of frequencies where they operate as stimulus feature detectors. We thus propose that the spectral content of structure-borne vibrations is extracted at the cortical level using a coding mechanism akin to the one operating in the auditory system for pitch discrimination of airborne sound. This sensory channel therefore appears to be fundamentally different from active skin- and whisker-based somatosensation.

## Results

### Vibration frequency selectivity in somatosensory cortex

To deliver substrate vibrations to a single forelimb, we trained head-restrained mice to place their forepaw on a piezoelectric stack actuator and reinforced continuous holding with liquid rewards (Fig. 1A,B, Movie S1, Methods). Pure sinusoidal vibrations (Fig. S1) of different frequencies (100 Hz to 800 Hz) were delivered pseudorandomly to the forepaw at variable inter-stimulus intervals. To measure the activity of neurons in the contralateral fS1, we expressed the fluorescent Ca^2+^ indicator GCaMP6f in layer 2/3 neurons using viral vectors and imaged the responses with two-photon microscopy through a chronic cranial window (Fig. 1C,D). The location of fS1 was identified by intrinsic signal imaging (Fig. 1D). We found that different vibration frequencies reliably evoked transient increases in fluorescence of a subset of imaged neurons (Fig. 1E,F). Since the size of fluorescence transients is positively correlated with the number of spikes in a burst (Chen et al., 2013) we took the fluorescence change relative to baseline (Δf/f_0_) as a proxy for spike rate. By observing the activity of individual neurons, it became evident that different stimulus frequencies did not evoke equal responses (Fig. 1F), suggesting that fS1 neurons carry information about vibration frequency in the number of spikes they fire.

**Figure 1.**
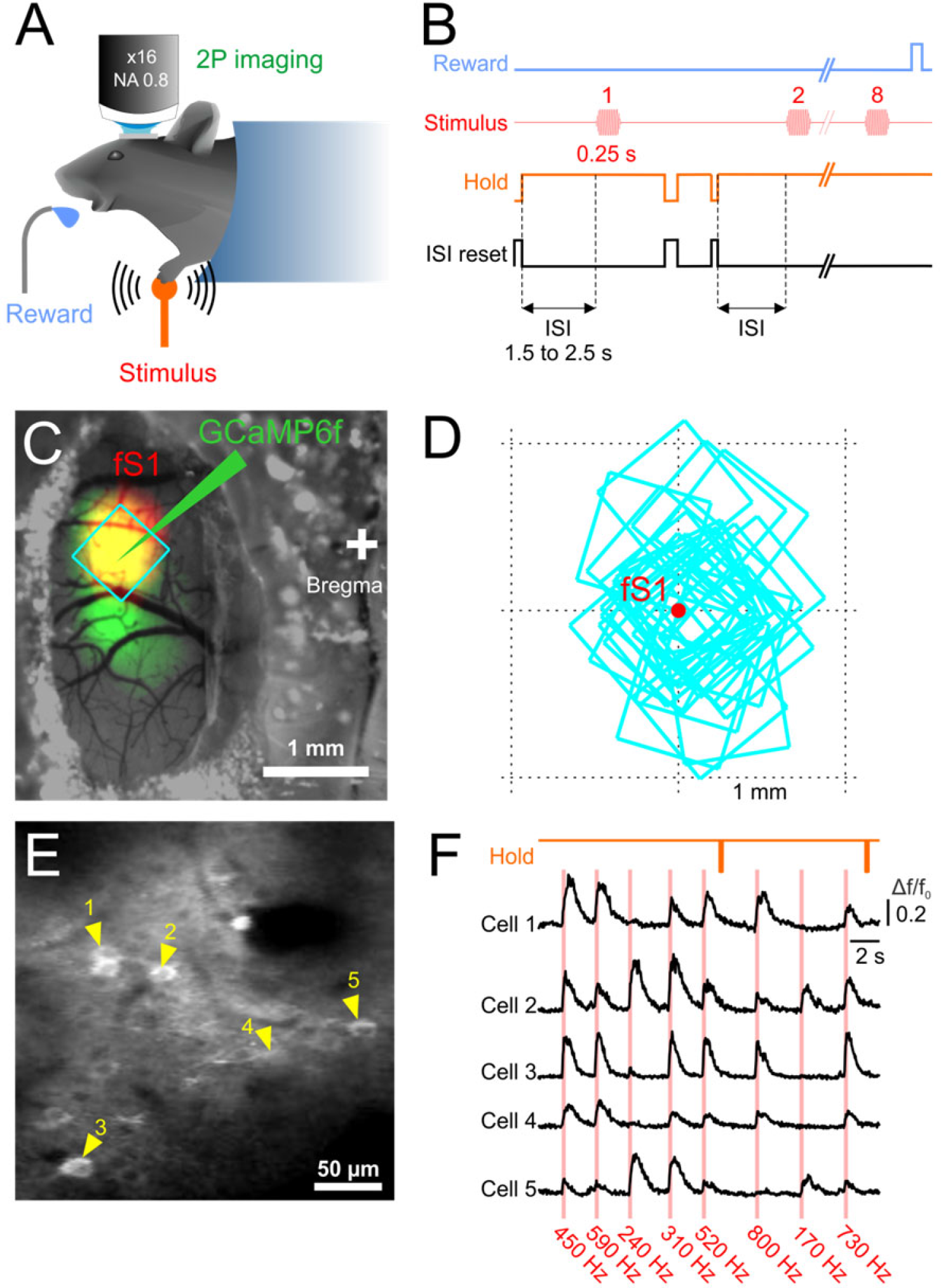
Two-photon Ca2_+_ imaging of forepaw vibration related activity in somatosensory cortex of awake mice. **(A)** Experimental setup schematic. **(B)** “Trial” timeline in which eight successive vibrotactile stimuli were delivered followed by reward. The variable inter-stimulus-interval (ISI) was reset upon forepaw release thereby ensuring contact with the stimulator at stimulus onset and incentivised continuous holding for higher reward rates. **(C)** Dorsal view of the cranial window showing the expression patterns of GCaMP6f (green), the intrinsic signal (red) of contralateral forepaw stimulation (fS1: forelimb somatosensory cortex) and the location of the imaging field-of-view (FOV, cyan square). **(D)** Locations of imaging FOVs (cyan squares) relative to the intrinsic signal center-of-mass of fS1 (red) of all experiments (19 mice, 42 FOVs). **(E)** Cropped in-vivo image of GCaMP6f expressing cells in fS1 with five neurons (yellow arrows) responsive to vibrotactile stimulation. **(F)** Imaged fluorescence changes in one “trial” of the five neurons shown in (E), in response to eight stimuli of different frequencies.

To explicitly test the latter hypothesis, we computed the stimulus-specific information (SSI) as a measure of frequency encoding by the evoked Δf/f_0_ responses (Butts, 2003; Butts and Goldman, 2006). The majority of responding neurons (>85% of 725 cells, 19 mice, 42 fields of view) were found to be informative about frequency (i.e. had responses with significant SSI, p<0.01, permutation test), which is in stark contrast to the 3% of neurons in the primate S1 responding to fingertip vibrations in the same frequency range (Harvey et al., 2013). To characterize their frequency dependent tuning, mean responses were fit with a descriptive function (Fig. 2A-H). Strikingly, ≈49% (303 of 621 informative cells) had identifiable tuning curve peaks (Fig. 2A-D,I) and thus had a preferred frequency of tactile vibration, which is analogous to a fundamental property of auditory cortex neurons: frequency selectivity (Cheung et al., 2001; Issa et al., 2014; Joachimsthaler et al., 2014). The remaining neurons were characterized by frequency dependent response increases-to-plateau or monotonic increases (Fig. 2E-H,I).

**Figure 2.**
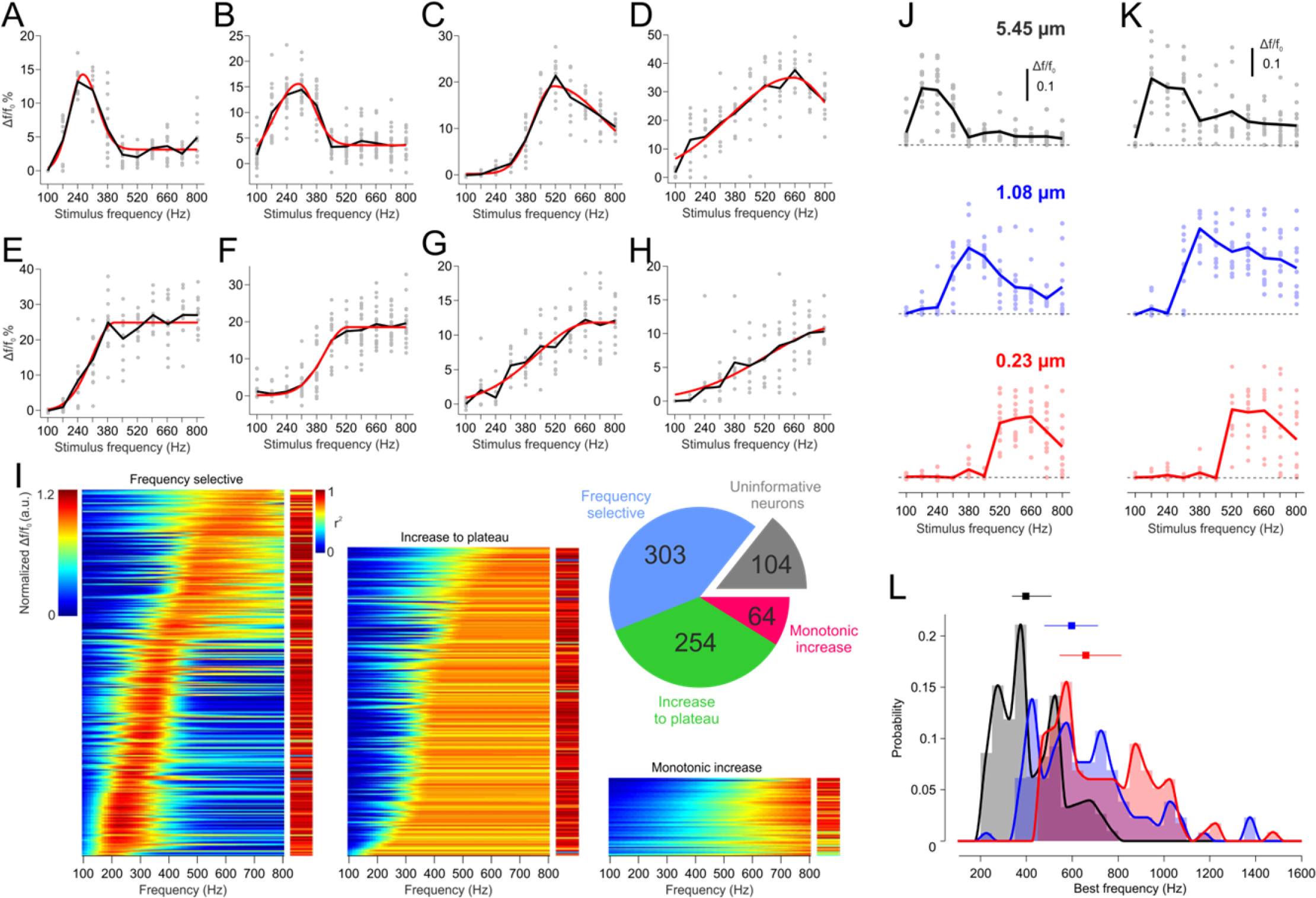
Frequency selective tuning to vibration frequency in mouse fS1. **(A-D)** Tuning curves of example frequency selective neurons (grey dots: individual stimulus responses, black lines: mean values, red lines: descriptive function fits). **(E-G)** Example neurons with increases-to-plateau tuning. **(H)** Example neuron with monotonic increase tuning. **(I)** Normalized tuning curve fits of all imaged neurons classified according to frequency selective, increases to plateau or monotonic increases with their respective goodness-of-fit measure (r^2^). The pie chart summarizes the number of neurons of each class, as well as those uninformative about stimulus frequency (i.e. with non-significant SSI).**(J,K)** Tuning curve changes of two example neurons as a function of vibration amplitude (dots: individual responses, lines: mean values). The black, blue and red tuning curves correspond to the same neuron tested at 5.45 μm, 1.08 μm and 0.23 μm vibration amplitudes, respectively. **(L)** Distribution histograms of best frequencies of all neurons with frequency selective tuning for the three tested amplitudes (5.45 μm: N=303 cells, 1.08 μm: N=130 cells, 4 0.23 μm: N=116 cells). Colors are as in (J,K) and lines are cubic interpolations of the histogram values. Squares with delimiters denote medians +/− quartiles.

We next assessed the effects of vibration amplitude on frequency tuning. As vibration amplitude was attenuated, a shift of the tuning curves toward higher frequencies was observed (Fig. 2J,K). Also, by reducing the actuator displacement, the range of frequencies that could be generated by the hardware was extended to 1990 Hz. As such, neurons characterized as having increases-to-plateau responses at the highest amplitude were found to also be frequency selective when tested over an extended frequency range with smaller displacements (Fig. S2). Based on the descriptive function fits, we identified neurons that are tuned to a preferred frequency at any of the three tested displacements. The distributions of their preferred values confirm that fS1 neurons best respond to higher frequencies at smaller stimulus amplitudes (Fig. 2L). The observed amplitude dependent changes in tuning (Fig. 2J,K, Fig. S2) are analogous to how frequency selective responses of auditory cortex neurons change as a function of sound level (Sutter, 2000).

### Frequency selective neurons operate as feature detectors

What might be the functional relevance of frequency selective tuning of fS1 neurons? Do these neurons signal the presence of a vibratory stimulus at their preferred frequency (i.e. operate as feature detectors) or is their role to differentiate stimuli based on their spectral content (i.e. perform frequency discrimination)? Modeling work has suggested that this question can be answered by analysing what the most informative region of the turning curve is (Butts and Goldman, 2006). On the curve’s flanks, where the slope is steepest, discrimination is best as small changes in stimulus value produce large changes in neural activity. At the curve peak, discrimination becomes worse as the slope is near zero, but the peak firing rate is most distinguishable from baseline activity and thus makes the preferred frequency distinct from all others. Using the SSI measure to identify the most informative stimuli, model neuron simulations demonstrate that as noise (i.e. the trial-to-trial variability in neural responses) increases, maximal SSI transitions from the tuning curve’s flank to its peak (Butts and Goldman, 2006). We therefore computed the SSI for all neurons with frequency selective tuning at any amplitude. The obtained results show that fS1 neurons convey maximum information at their preferred frequency (Fig. 3) and therefore operate, just like neurons in auditory cortex (Montgomery and Wehr, 2010), in a high-noise regime as feature detectors.

**Figure 3.**
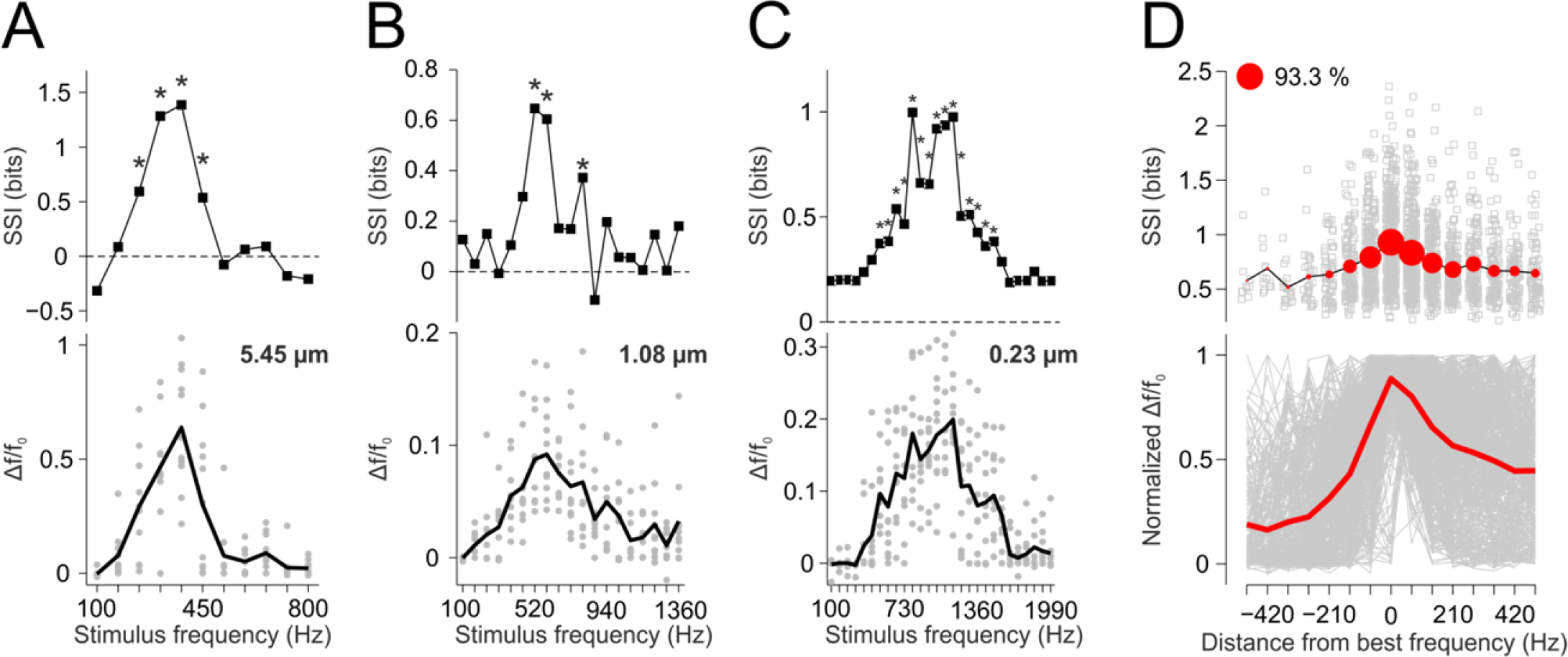
Maximum information is conveyed at the tuning curve peaks. **(A-C)** Frequency tuning of three representative neurons (bottom) at indicated vibration amplitudes and their respective bias-corrected SSI (top) *: p<0.01, permutation test (n=1999 permutations). **(D)** Normalized tuning curves (grey) and the average (red) of all frequency selective neurons at any amplitude (N=549 cells) aligned to their best frequency (bottom) and the corresponding significant SSI values (grey squares), their mean (black line) and their proportions (red circle sizes) relative to the total number of neurons (top).

### fS1 neuronal spikes are not cyclically entrained by forepaw vibrations

At the periphery, each cycle of a sinusoidal oscillation applied to the skin typically evokes one or more phase-locked spikes in mechanoreceptor afferents (Mackevicius et al., 2012; Talbot et al., 1968). It follows that spike rate monotonically increases with vibration frequency up to the afferent’s best frequency. At the cortical level, in S1 neurons representing the primate hand (Harvey et al., 2013; Mountcastle et al., 1969) and rodent whiskers (Ewert et al., 2008), the cycle-to-cycle tracking of the vibration is lost but phase-locked entrainment of spikes persists. Spectral information is therefore only encoded in their temporal spiking patterns and not anymore in firing rates. Given our finding that frequencies are encoded in firing rates in fS1 when substrate vibrations are applied to the forelimb, we wondered whether phase-locked spiking also exists. To address this, we measured stimulus-related spiking with juxtacellular (37 units, 6 mice) or extracellular (17 units, 6 mice) electrode recordings using identical conditions as in imaging experiments (Fig. 2A and Fig. S3A). We found frequency dependent increases (43 units, Fig. S3B-G), but also decreases (11 units, Fig. S3H,I) in spike rates, with a similar heterogeneity of tuning curves as inferred from the imaged calcium fluorescence. For the rate increasing units, we measured whether a preferred phase of spike occurrence in the stimulus cycle exists (Fig. 4B). Phase distributions did not reveal significant phase-locked spiking (Fig. 4C) at any frequency (Rayleigh test for non-uniformity of circular data, p<0.01 for 1 out of 43 units at most). To control for possible variability in onset delays across stimulus repetitions, we also computed inter-spike-intervals whose distribution should peak at integer multiples of the stimulus period in the case of phase-locked spiking. At any stimulus frequency, the null hypothesis that standardized inter-spike-intervals (see Methods) come from a uniform distribution (Fig. 4C) could be rejected for at most 1 out of the 43 units (χ^2^ test, p<0.01). We therefore speculate that a transformation of neural coding, analogous to that in the auditory system (Saal et al., 2016), occurs between peripheral and central processing of stimuli vibrating the mouse forelimb, where phase-locked spiking progressively dissipates along the ascending pathway and turns into a pure rate code in the cortex.

**Figure 4.**
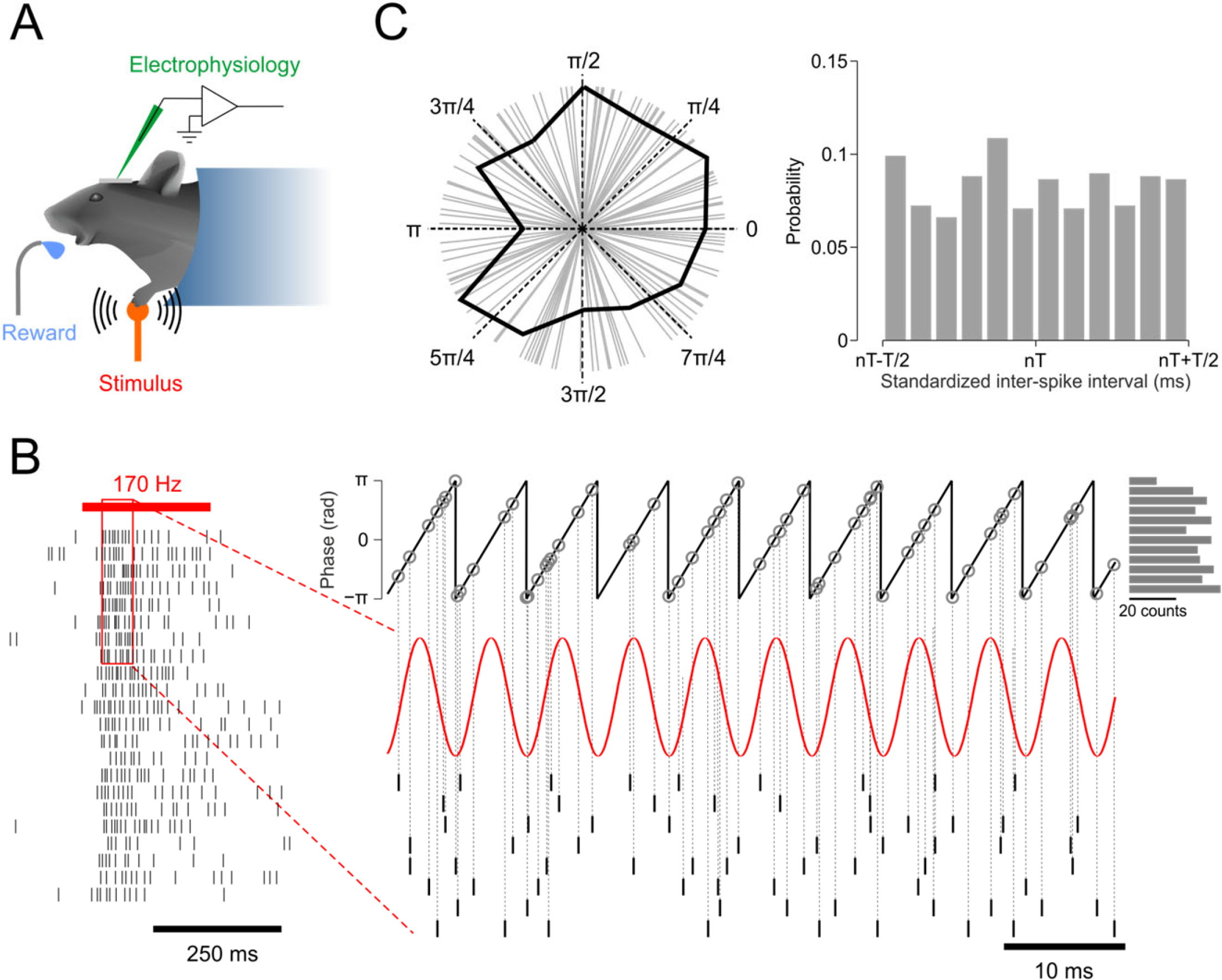
The spikes of fS1 neurons are not cyclically entrained by the sinusoidal vibration. **(A)** Experimental setup schematic. **(B)** Spike raster plot of a neuron responding to a 170 Hz vibration. To test for phase locked spiking, the distribution of phases of the stimulus wave at times of spike occurrences was computed as depicted, for each responding neuron and all frequencies. **(C)** Left: distribution of stimulus phases at spike times for a representative neuron at its best frequency (grey lines: phases of individual spikes, black contour: circular phase histogram). Rayleigh test for non-uniformity of circular data: p=0.395. Right: histogram of standardized inter-spike-intervals of the same example neuron. χ^2^ test for uniformity: p=0.1.

### Responses are not driven by auditory cues

The striking similarity to response properties of auditory neurons raises the possibility that airborne sounds, produced by the piezoelectric stack actuator generating the vibratory stimulus, drive activity in fS1 neurons. Alternatively, the substrate vibrations might propagate through the bones to the inner ear (Rado et al., 1998; Reuter et al., 1998). Auditory tones are known to alter the perception of tactile stimuli in a frequency-specific manner (Crommett et al., 2017; Yau et al., 2009) and might hence indeed drive activity in the somatosensory cortex. To test for these possibilities, we placed the forepaw away from the stimulator or saturated the ear with white noise, and compared the responses to those evoked by normal control conditions in the same experimental sessions. With the forepaw placed off the stimulator (113 cells, 9 mice), sound alone could not activate, at any frequency, the neurons normally responsive to vibratory stimulation. Masking hypothetical inner ear vibrations with white noise (128 cells, 9 mice) caused only small changes in the evoked responses (Fig. S4), probably due to the averseness of the loud sound. These results demonstrate that the auditory system is not implicated in the frequency tuning of fS1 neurons. Additional experiments in which mice were trained to perform reach-to-grasp forelimb movements (Movie S2) allowed us to exclude hypothetical stimulus-locked motor events such as muscle twitches, increases in stiffness or proprioceptive inputs (Fig. S5).

### Pacinian corpuscles are absent in the glabrous skin of the mouse forepaw

In mammals, high frequency vibrations are transduced by a subtype of rapidly adapting mechanoreceptors, the Pacinian corpuscles (PCs) (Abraira and Ginty, 2013). PCs can be found in many body locations near joints, ligaments, tendons and bones (Bell et al., 1994; Zelena, 1994), but they most prominently populate the glabrous skin of fingertips and palm of the primate hand (Kumamoto et al., 1993a). Is it possible that the cortical coding scheme for substrate vibration frequency, reported here, originates from the recruitment of a functional network of PCs different to that recruited in the primate hand by local cutaneous vibrations?

To address this question, we performed a detailed histological analysis of PC distribution in the mouse forelimb. PCs were identified in hematoxylin and eosin stained sections of the distal limb based on their stereotypical lamellar morphology (Kumamoto et al., 1993a; Kumamoto et al., 1993b). Unlike in the primate hand, we were unable to find PCs in the dermis of the glabrous forepaw skin (Fig. S6), which confirms previous observations (Fleming and Luo, 2013). However, PCs were systematically found in deep tissue next to bones in the fingers and palm, but also more proximally next to the radius and ulna (Fig. 5). This finding suggests that the cortical responses reported here originate not from the activation of dermal but rather of a population of deep PCs.

**Figure 5.**
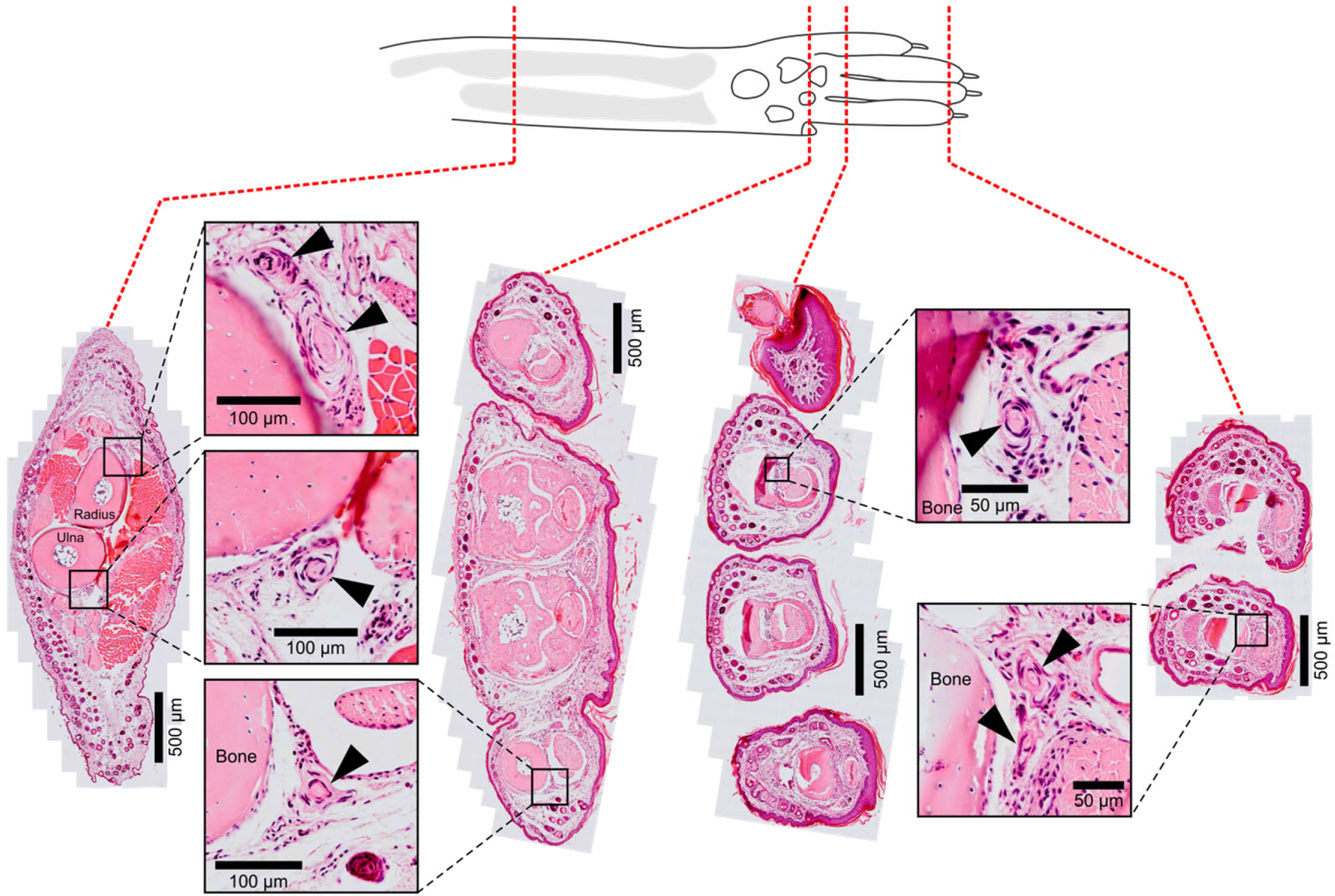
Pacinian corpuscles are located subcutaneously in the mouse forelimb. Four transverse 5 μm sections (haematoxylin and eosin stained) of a mouse forelimb show Pacinian corpuscles (black arrows) located deep in subcutaneous tissue next to the bones of fingers and the radius/ulna. The Pacinian corpuscles were never found in the skin dermis.

### Responses are specific to mechanoreceptor activation in the distal forelimb

To causally link the frequency tuning observed in fS1 to the activation of mechanoreceptors in the distal forelimb we conducted two additional experiments. First, we blocked nerve transmission from the forepaw by injecting lidocaine at the wrist level and compared the evoked activity to the pre-injection control period (83 cells, 8 mice). The induced changes were compared to the effects of saline injections performed either the following or previous day with the same neurons. The significantly higher reduction of the evoked responses with lidocaine (Fig. 6A) suggests the involvement of mechanoreceptors in the wrist/forepaw in driving the imaged neurons. However, the nerve block did not uniformly reduce responses across the entire frequency range (Fig. 6B-D). This frequency-dependent effect might be analogous to tuning curve changes observed when vibration intensity is reduced (Fig. 2J-L and Fig. S2). Alternatively, the substrate vibrations are likely to propagate to and activate mechanoreceptors in other parts of the forelimb (Fig. S5) or even in more distant organs (Bell et al., 1994; Zelena, 1994). The residual responses after lidocaine injection might thus reflect their frequency selective contribution that converges together with that of mechanoreceptors in the wrist/forepaw on the same cortical fS1 neurons.

**Figure 6.**
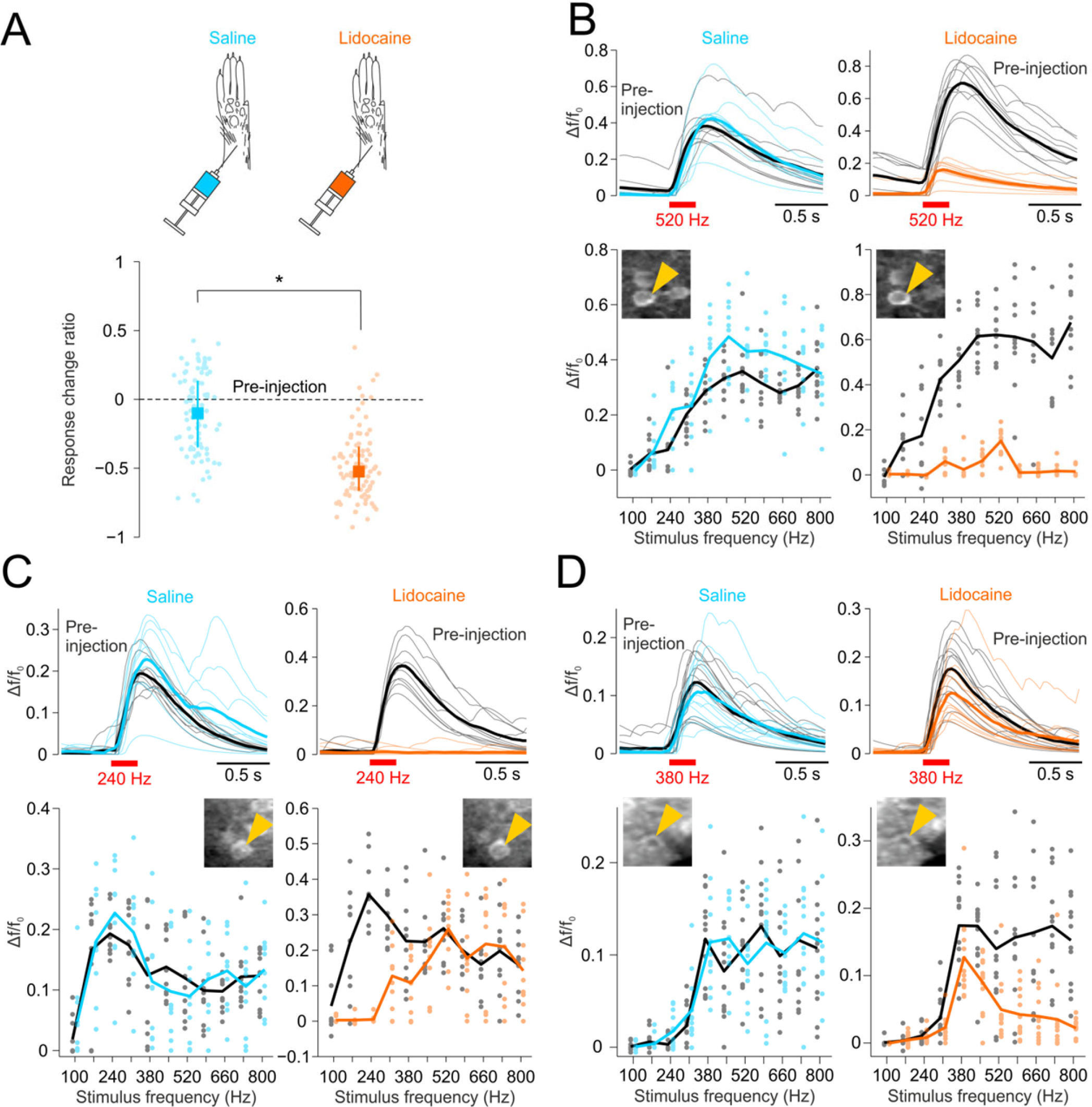
Response specificity to forelimb stimulation tested with peripheral nerve blocking. **(A)** To test if the imaged neural responses were specific to vibrotactile stimulation of the forepaw, nerve transmission was blocked by lidocaine injection at the wrist level. Equal volume (50 μL) saline injection was used as control. For each neuron, the response change ratio (median +/− quartiles, n=83 cells, 8 mice) relative to pre-injection activity (zero level) was averaged across all frequencies with significant responses in the pre-injection period. *: p<10^−12^, Wilcoxon signed rank test. **(B)** Representative neuron showing supressed responses across all tested frequencies after lidocaine (orange) but not saline (cyan) injection compared to pre-injection levels (black). Top: denoised Δf/f_0_ responses evoked by 520 Hz vibrations (thin lines: individual stimulus responses, thick lines: mean response). Bottom: frequency tuning curves (dots: individual responses, lines: mean response). Insets: cropped two-photon images showing that the same neuron was analyzed in both experiments (yellow arrow). **(C,D)** Same data as in (B) for two representative neurons showing supressed responses in a restricted range of frequencies only.

In a second set of experiments, we expressed the optogenetic actuator Channelrhodposin-2 selectively in a subset of mechanoreceptors associated with rapidly adapting afferents. Pacinian and Meissner corpuscles and their innervating neurons were targeted using a double-transgenic strategy (ETV1-Cre x ChR2-EYFP, see Methods). Two-photon Ca^2+^ imaging of fS1 activity was performed while either vibrotactile, optogenetic (Movies S3,S4) or single-point touch (see Methods) stimuli were delivered to the contralateral forepaw. Pulsed illumination of the forepaw with an optic fiber evoked highly reliable Δf/f_0_ increases in a subset of cortical fS1 neurons (20 cells, 2 mice, 3 fields of view) (Fig. 7A-C and Fig. S7A). We grouped trials with similar paw placement relative to the optic fiber by applying a dimensionality reduction technique, t-distributed stochastic neighborhood embedding (t-SNE) (van der Maaten and Hinton, 2008a), to forepaw images at the time of stimulus onset (Fig. S7B). This classification revealed that the fS1 neurons were preferentially activated by illumination of specific locations on the forepaw (Fig. 7A-C and Fig. S7C-E). Some of the same neurons also responded to either vibration (Fig. 7A) or single-point touch (Fig. 7B,C). In sum, our vibratory stimuli were found to drive cortical neurons intermingled and partially overlapping with those responding to touch of the forepaw and to selective optogenetic activation of its mechanoreceptor afferents (Fig. 7D). This result supports the assertion that the frequency selective fS1 neurons, reported here, reflect the processing of substrate vibrations acting upon mechanoreceptors of the distal forelimb.

**Figure 7.**
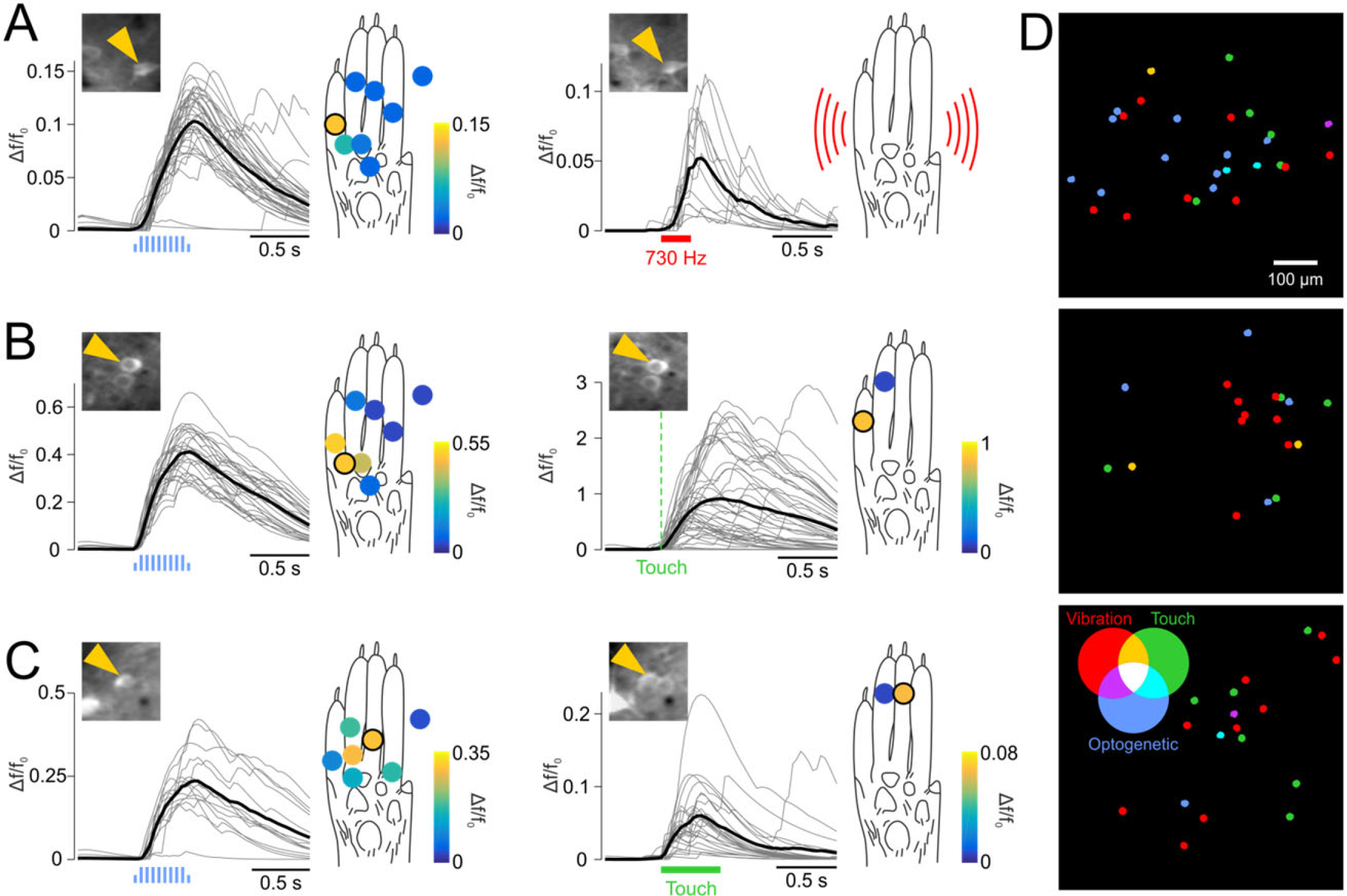
Response specificity to forelimb stimulation tested with optogenetic tagging. **(A)** Example neuron in a double transgenic mouse activated by both optical stimulation (left) and vibration (right) of the forepaw. Filled circles indicate different stimulation locations and are color coded according to the average Δf/f_0_ response. The depicted denoised Δf/f_0_ response traces correspond to the stimulus location with the black outline and the 730 Hz vibration (thin lines: individual stimulus responses, thick lines: mean response). Insets: cropped two-photon images showing that the same neuron was analyzed in both experiments (yellow arrow). **(B,C)** Same data as in (A) for two example neurons activated by both the optical stimulation and either manual (B) or automatized (C) single point touch. The black Δf/f_0_ traces correspond to touch locations (filled black circles) from which significant responses could be evoked. The orange traces and symbols correspond to control touch locations. **(D)** The distribution of neurons within three imaged FOVs (2 mice) with significant responses to either optogenetic, vibration or touch stimuli, or any combination of the three as indicated by the color code.

## Discussion

We have shown that vibrating the mouse forelimb with pure sinusoidal displacements evokes auditory-like responses in somatosensory cortex neurons: frequency selective spike rates and absence of cyclical spike entrainment. Both of these findings are at odds with previous reports of how S1 neurons code for the frequency of tactile vibration applied to a single point on the skin (e.g. at fingertips) (Harvey et al., 2013; Mountcastle et al., 1969). How can this discrepancy be explained?

Localized oscillatory indentations of the skin surface constitute ecologically relevant vibrotactile sensations for primates, which might occur during active tactile exploration. The associated sensation is most likely conveyed to the central nervous system by dermal PCs that densely populate the primate fingertips (Kumamoto et al., 1993a). On the contrary, we found no evidence that dermal PCs exist in the mouse forepaw (Fig. 5 and Fig. S6), which has been previously reported (Fleming and Luo, 2013). This finding might also support the view that rodents principally use their whiskers rather than their paws for active tactile exploration (Diamond et al., 2008) (but see (Manita et al., 2015; Maricich et al., 2012)). Forepaw-based vibration sensation in rodents might thus take the role of passive “listening” through the substrate as the communication medium (Hill, 2008). The activation of deep, subcutaneous PCs populating the distal forelimb at large might synergistically participate in processing the vibratory stimuli. The associated activity in the multiple afferent channels might be individually filtered and eventually converge onto cortical neurons with frequency selective spike rates. The frequency-dependent effect of our pharmacological experiments (Fig. 6) and the absence of a one-to-one correspondence between neurons driven by selective afferent activation and forepaw vibration (Fig. 7D) are consistent with this hypothesis. On the other hand, localized skin vibrations in primate fingertips drive cortical neurons with firing properties corresponding more closely to those of individual dermal PC afferents (Mackevicius et al., 2012; Talbot et al., 1968).

Besides the difference in receptor distribution found between the mouse and primate, we can ask more generally: do the two different spectral information coding schemes (rate code vs. cyclical entrainment) have different functional implications? We speculate that they might reflect a fundamental difference between active and passive sensing. Active somatosensory systems need to take the kinematics of the active motion into account when extracting low-level stimulus features. Such timing dependent sensory-motor integration might therefore explain temporally precise coding in the cortex, as described by cyclical spike entrainment for example (Ewert et al., 2008; Harvey et al., 2013; Mountcastle et al., 1969). Instead, cortical feature-selective rate coding in the primate fingertip or the rodent whisker systems have most likely evolved to identify macroscopic object features like surface texture (Garion et al., 2014; Isett et al., 2018) or edge orientation (Bensmaia et al., 2008), independently of motion kinematics. On the other hand, passive sensory systems, such as hearing or sensing substrate vibrations do not rely on active motion and have most likely evolved to directly identify and discriminate lower level physical parameters like frequency or pitch using a tuned rate code (Eggermont, 2001). This distinction is directly supported by psychophysical experiments comparing how humans perceive the spectral information of sound and skin vibrations (Formby et al., 1992). However, both coding schemes are probably highly overlapping and might even use the information of the same receptors, yet via segregated circuits. Future experiments will be necessary to investigate if such parallel coding systems exist, for example at the level of the primate hand.

Interestingly, we found that the responses to forelimb vibrations described here not only share many features with the auditory system, the frequency range represented (100 Hz to 1.5 kHz, Fig. 2L) is also surprisingly complementary to the auditory hearing range of the mouse (4kHz to 50kHz) (Koay et al., 2002). The passive sensing of substrate vibrations might thus be a natural extension of the mouse auditory system to the lower, inaudible frequency range. Actually, it has been proposed that communication via airborne sounds might have evolved from the more ancient precursor modality based on substrate-borne vibration signaling (Hill, 2008). Many insect species communicate exclusively by emitting and sensing substrate vibrations (Cocroft and Rodríguez, 2005) while in others, the same sensory organ, such as the Johnston’s organ in drosophila, is used to detect both sound and touch (Azevedo and Wilson, 2017). Vestiges of this modality seem to be still present in rodents, given that Ehrenberg’s mole-rats vibrate their subterranean tunnels to communicate with conspecifics (Heth et al., 1987; Rado et al., 1987), and might explain the parallels that we found between frequency coding in auditory and somatosensory systems.

In conclusion, our work reveals a novel and robust neural representation of vibration frequency in the somatosensory cortex. The remaining challenge is to understand the mechanisms by which this cortical representation emerges from activations of peripheral mechanoreceptors along the ascending pathway. Similarly to recent experiments of primary afferents in the whisker system (Severson et al., 2017), we developed an optogenetic tagging strategy to selectively activate rapidly adapting mechanoreceptor afferents in the forepaw. We were furthermore able to identify the neurons that this peripheral optogenetic activation drives in the cortex and compare them to those activated by natural tactile stimuli (Fig. 7). These experiments therefore pave the way for resolving the above posed challenge and, more generally, for future studies to illuminate the functional contribution of a given sensory receptor to cortical processing of sensory information.

## Methods

### Animals

Imaging and electrophysiology experiments were conducted with C57BL/6 (Charles River Laboratory) mice. Forepaw optogentic stimulation experiments were conducted with double-transgenic mice generated by mating homozygote Ai32 males carrying a floxed ChR2-EYFP fusion gene inserted in the *Gt*(*ROSA*)*26Sor* locus in a C57BL/6 strain (Madisen et al., 2012) (Jackson Laboratory; stock no. 012569) with heterozygote ER81/Etv1-CreER females expressing the CreER^T2^ fusion protein from the *ER81/Etv1* promoter elements (Taniguchi et al., 2011) (Jackson Laboratory; stock no. 013048). As such, Cre-mediated recombination results in an expression of the floxed ChR2-EYFP sequence in the ER81/Etv1-expressing cells of the offspring. ER81/Etv1 is expressed in Pacinian and Meissner corpuscles, glabrous skin mechanoreceptors associated with rapidly adapting afferents, as well as in their innervating neurons (Fleming et al., 2016). To induce CreER-based recombination, double-transgenic offspring (10 week old) were administered one daily dose of a 100 μL tamoxifen (T5648, Sigma-Aldrich) solution (20 mg/mL) dissolved in sunflower oil (C8267, Sigma-Aldrich) for five consecutive days by peritoneal injection. Mice were housed in an animal facility, maintained on a 12:12 light/dark cycle and were placed under a water restriction regime (1 ml/day) post-surgery. The experiments were performed during the light phase of the cycle. The animals did not undergo any previous surgery, drug administration or experiments and were housed in groups of maximum 5 animals per cage. All procedures complied with and were approved by the Institutional Animal Care and Use Committee of the University of Geneva and Geneva veterinary offices.

### Surgery

12 to 15 week old mice were surgically prepared for head-fixed two-photon Ca^2+^ imaging. Surgeries were conducted under isoflurane anaesthesia (1.5 to 2%) and additional analgesic (0.1 mg/kg buprenorphine intramuscular (i.m.)), local anaesthetic (75 μL 1% lidocaine subcutaneous (s.c.) under the scalp) and anti-inflammatory drugs (2.5 mg/kg dexamethasone i.m. and 5 mg/kg carprofen s.c.) were administered. Mice were fixed on a bite bar with a snout clamp and rested on top of a heating pad. The scalp was cut over the midline between the ears and eyes. A custom made titanium head bar was fixed on the skull with a cyanoacrylate adhesive (ergo 5011, IBZ Industrie) and dental cement to allow for subsequent head fixation. A craniotomy was performed over the left frontal cortex and five viral injections (30 to 50 nL each) were administered with pulled and beveled glass pipettes (≈25 μm tip diameter) at a an approximate rate of 20 nL/min into the forepaw representation of the primary somatosensory cortex (2.25 mm lateral and from −0.25 mm to 0.75 mm in 0.25 mm steps anterior to bregma). The injections consisted of AAV1.Syn.GCaMP6f.WPRE.SV40 (1:10 dilution with 0.2% FastGreen in sterile saline, virus stock titer 6.932 × 10^13^) (University of Pennsylvania, lot. no. CS0939) at a depth of 350 μm corresponding to cortical layers 2/3. After each injection completion, the pipette was left in place for at least another 10 min before being retracted. The cortical surface was rinsed for 1 to 2 min with dexamethasone (0.03%) and two hand-cut glass coverslips (150 μm thickness) matched to the shape of the craniotomy were glued together with optical adhesive (NOA 61, Norland Products) and the lower one was placed on top of the cortex. The upper glass, which was larger than the craniotomy, was glued to the skull with cyanoacrylate adhesive and secured with dental cement. The correspondence between the injection sites and the sensory forepaw representation was confirmed post-operatively with intrinsic signal imaging in each mouse. After a seven day recovery period, mice were placed under the water restriction regime and experiments began on the 15^th^ day after surgery.

### Intrinsic signal imaging

Mice were head-fixed and placed on a heated platform under light isoflurane anaesthesia (0. 25 to 0.75%). Their right forepaw rested on a metal rod (2 mm diameter) mounted on a galvanometric actuator (model. no. 000-G120D, GSI Lumonics). Ten 1 s vibrotactile stimuli consisting of a 100 Hz sinusoidal vibration were delivered to the forepaw with a 10 s inter stimulus interval. The cranial window was illuminated by a collimated red light LED (635 nm peak, ACULED VHL, part. no. E001947, Perkin Elmer) and imaged with a CCD camera at 10 fps with a 256 by 332 pixels resolution (Retiga-2000R, Q Imaging). The average difference image between the stimulation period and a 1.5 s baseline period was processed by a 50 by 50 pixels spatial averaging filter and subsequently smoothed by a 5 by 5 pixels Gaussian low pass filter with 0.5 pixels standard deviation. Stimulus presentation and image acquisition were controlled by Ephus software (Scanimage.org). The expression pattern of GCaMP6f was imaged by illuminating the cranial window with a collimated blue LED light (455 nm peak, ACULED VHL, part. no. E001947, Perkin Elmer) bandpass filtered between 465 and 495 nm (EX465-495 FITC, Nikon). Before detection, the emitted green light was bandpass filtered between 515 and 555 nm (BA515-555 FITC, Nikon).

### Two-photon Ca^2+^ imaging

Imaging was performed with a custom built two-photon microscope (MIMMS, www.openwiki.janelia.org) controlled by Scanimage 5.1 (Vidrio Technologies) using a 16x 0.8 NA objective (Nikon) and with excitation wavelength at 940 nm (Ultra II, tunable Ti:Sapphire laser, Coherent). 512 by 512 pixel images covering 626 by 665 µm of cortex were acquired at 29.82 Hz using bidirectional scanning with a resonant scanner system (Thorlabs). The power was modulated with pockels cell (350-80-LA-02, Conoptics) and calibrated with a photodiode (Thorlabs). The primary mirror for imaging was a custom polychroic (Chroma, zt470/561/nir-trans) transmitting the infrared light, while reflecting the green. Before detection, the remaining infrared light was filtered with a colored glass band pass filter (BG39, Chroma) while blue light used for optogenetic forepaw stimulation was removed with a short pass filter (CG475, Chroma). Images were acquired using photo multiplier tubes (H11706P-40 SEL, Hamamatsu) and written in 16 bit format to disk. The trial start, reward and stimulus onset TTL pulses issued by the behavior PC were received as auxiliary triggers and their timestamps saved in the headers of the acquired images. The saved timestamps were used for temporal alignment of Ca^2+^ traces with behavioral variables during post-hoc analysis.

### Electrophysiology

In vivo recordings were performed in awake head-fixed mice (12 to 15 weeks old) and targeted to fS1. A small craniotomy was drilled over fS1 in mice surgically implanted with a head bar as described above. The electrode was an Ag/AgCl wire inserted into an internal solution filled glass pipette. The internal solution contained (in mM) K-gluconate 134, KCl 6, HEPES 10, NaCl 4, Mg_2_-ATP 4, Tris_2_-GTP 0.3, Na-phosphocreatine 14, pH 7.25. The pulled glass pipettes had a tip-resistance of 6 to 12 MΩ. A second reference Ag/AgCl electrode connected to the headstage (CV-7B, Axon Instruments) was placed inside the recording chamber filled with saline. Positive pressure was applied as the pipette was lowered until 100 µm below the pial surface and then advanced in 2 μm steps. When action potentials were encountered, their wave shape had an initial negative phase. If upon further electrode advancement, the initial phase flipped to positive and action potential amplitude increased to many orders of magnitude above baseline, the recording was considered to be juxtacellular, and extracellular otherwise. All recordings were done in current clamp mode. The signal was amplified and filtered (0.1 Hz to 10 kHz) at the headstage and acquired at 30 kHz (PCI-MIO-16E-4, National Instruments) using custom Matlab (Mathworks) routines. Trial start and stimulus onset triggers were saved in parallel on separate channels and used for post-hoc alignment of recorded spikes.

### Pharmacology

To block neural transmission from the forepaw, mice were briefly anaesthetized with isoflurane (3%) and 50 μL of Lidocaine (1%) was injected subcutaneously at level of the palmar wrist. Mice were subsequently head-fixed under the two-photon microscope and allowed to recover from anaesthesia. Vibrotactile responses were imaged from 12.6 ± 2.6 min to 17.7 ± 3.1 min post-injection and compared to responses imaged ≈10 min pre-injection. The same procedure was repeated on either the previous or following day with 50 µL of saline with responses being imaged from 13.2 ± 3.5 min to 18.3 ± 3.7 min post injection.

### Histology

After hair removal and PFA 4% intracardiac perfusion, mouse forelimbs were sectioned at the elbow. The samples were post-fixed in PFA 4% overnight and washed in PBS. The sectioned forelimbs were decalcified by immersion in EDTA-Citronensäure-Puffer pH 7.5 (ProTaqs, BIOCYC GmbH & Co.) for 15 days at room temperature. The decalcifying agent was changed once per week and decalcification was deemed complete by manually testing the bone resistance of each sample. 5 μm thick transverse sections were taken between elbow and fingertips and embedded in paraffin. The sections were stained for eosin and hematoxylin using a standard protocol.

### Vibrotactile stimuli

The vibrotactile stimuli were generated by a piezoelectric stack actuator (P-841.3, Physik Instrumente) equipped with a strain gauge feedback sensor. A steel rod (2 mm diameter) on which mice placed their forepaw was mounted on the actuator. The actuator and sensor controllers (E-618.1 and E-509.S1, Physik Instrumente) operated in closed loop thereby counteracting any force the mouse applied. Pure sinusoids (250 ms duration, 25 ms linear onset/offset ramps) of 11 different frequencies (100 to 800 Hz in 70 Hz increments) calibrated to produce a 5.45 µm displacement amplitude were sampled at 10 kHz (USB-6353, National Instruments) and fed to the actuator controller. The sensor measurements were continuously acquired and recalibration of motor commands was regularly performed for the stimuli to remain highly consistent. The spectrum of the acquired sensor measurements was analyzed post-hoc to ensure the integrity of their frequency content (Fig. S1). The 800 Hz upper bound was imposed by the controller’s bandwidth for the 5.45 µm displacement. Two additional attenuated displacement amplitudes were tested with an extended frequency range: 1.08 µm (100 to 1360 Hz) and 0.23 µm (100 to 1990 Hz).

### Optogenetic stimuli

Optogenetic stimuli were generated by blue light illumination (472 nm, 50 mW; Obis FP 473, Coherent) out of a fiber (3.5 μm core diameter, 0.045 NA). The stimulus was a 20 Hz train of square pulses (500 ms duration, 25 ms pulse length) with the first and last pulses of the train set to half power (25 mW). The fiber was inserted and fixed in the center of a 3D printed holder (3.5 mm diameter, 7.5 mm length) on which the mice placed their forepaw. The stimuli were delivered using the same trial structure as with vibrotactile stimuli.

### Touch stimuli

Touch stimuli were delivered either manually or in an automatized manner with a custom built device. Manual touches of different parts of the forepaw were gentle skin indentations using a 24 gauge needle. For automatized indentations, a metal rod (0.5 mm diameter) was mounted on a miniature 8 Ω speaker (32KC08-1, Veco Vansonic) and guided through a tube (3 mm outer diameter) that was inserted and fixed in the center of the 3D printed paw holder (see above). Touches were generated by relaying a 3 V signal to the speaker for 0.5 s and delivered using the same trial structure as with vibrotactile stimuli.

### Behavioral procedures

Mice sat head-fixed in a tube (25 mm inner diameter) and were trained to place the forepaw contralateral to the imaged cortical site on the stimulation device. The ipsilateral forelimb was blocked from protruding outside the tube. On every trial, eight successive stimuli were delivered with a randomly varied inter-stimulus-interval (1.5 s to 2.5 s) followed by a water reward. After reward consumption, a new trial started. Paw placement was monitored with a digital fiberoptic sensor (FX-301, Panasonic). Paw removal resulted in resetting the inter-stimulus-interval thereby enforcing that mice hold the stimulator at every stimulus onset. Frequent paw removals therefore prolonged trial duration and delayed reward delivery, incentivising mice to continuously hold the stimulator to achieve higher reward rates. Mice were trained ≈1 h/day for 7 to 10 days before the start of Ca^2+^ imaging experiments. The different vibrotactile frequencies were randomly sampled without replacement and at least 10 repetitions of each frequency were tested in a given imaging session.

The described experimental design enforced that mice hold the vibration device upon stimulus presentation. Evoked responses could therefore not have been related to overt forelimb movements. However, stimulus-triggered muscle twitches, increases in stiffness or proprioceptive inputs cannot be excluded. We reasoned that neurons associated with these covert motor acts would be activated by reach-to-grasp forelimb movements. Accordingly, as a control experiment, we modified the task by training mice to reach for the water reward instead of delivering it at the snout following the series of eight stimulus presentations. For this purpose, the water drop was presented at the opening of a descending metal tube placed at reaching distance and mice executed forelimb reaches to grasp and consume the droplet (Movie S2). Mice were trained ≈1 h/day for 10 to 14 days on this task before the start of Ca^2+^ imaging experiments.

All behavior was controlled with real-time routines running on Linux (Bcontrol, brodylab.princeton.edu/bcontrol) and interfaced with Matlab (Mathworks) running on a separate PC.

## Two-photon image analysis

### Movement correction

To correct for lateral movements, a custom MATLAB registration algorithm was used to align each image to a template taken as the average image of a 30 s resting baseline period recorded at the start of each session. The cross-correlation was computed between each image and the template by multiplying the two-dimensional discrete Fourier transform of one with the complex conjugate of the Fourier transform of the other and taking the inverse Fourier transform of the product. The row and column location of the peak cross-correlation value was taken as the vertical and horizontal shift, respectively. 10% of each image was cropped at the boundaries for the purposes of this computation.

### Region of interest and Ca^2+^ activity generation

Using the session mean and variance images, soma centers of active neurons with clearly identifiable morphologies were manually initialized. Regions of interest of individual neurons and background were then identified as spatial footprints using the constrained nonnegative matrix factorization method (Pnevmatikakis et al., 2016) from the ca_source_extraction Matlab toolbox (gitgub.com/epnev/ca_source_extraction). This algorithm also provided the time-varying calcium activity of each neuron and background/neuropil as well as their time-varying baseline fluorescence. The two signals were used to compute Δf/f_0_ traces used for analysis with the modeled background activity removed. A denoised Δf/f_0_ trace of each cell was inferred with a constrained deconvolution approach.

## Data analysis

### Significant responses

The stimulus-evoked Δf/f_0_ response of a given neuron was computed as the difference between the maximum Δf/f_0_ value post stimulus onset (between 0 and 0.6 s relative to onset) and the mean Δf/f_0_ value pre-stimulus onset (between 0 s and −0.25 s relative to onset). This computation was repeated 1999 times for randomly shifted stimulus onset times across the neuron’s activity trace of the entire session. In this manner, 1999 chance measures of the average Δf/f_0_ response could be compared to the true Δf/f_0_ response means for each vibration frequency. If the true mean was more extreme than the upper 99^th^ percentile of the chance values distribution, the neuron was deemed to be responsive to that particular stimulus frequency. This procedure constitutes a randomization test at significance level p<0.01.

### Stimulus specific information

As a measure of frequency (ƒ) encoding by a neuron’s evoked Δf/f_0_ responses (*r*) we calculated the stimulus specific information (*SSI*) according to the method of Butts and Goldman (Butts, 2003; Butts and Goldman, 2006). The responses were first binned into one of 11 uniformly distributed values. We then computed the entropy of the stimulus ensemble

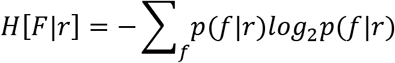

and the entropy of the stimulus distribution conditional on a particular binned response value

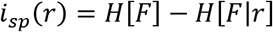

*p*(ƒ) is the fraction of stimuli with frequency *f* and *p*(*f*|r) is, for a particular response *r*, the fraction of stimuli with frequency f.

The specific information of a response

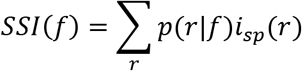

represents the amount learned about the stimulus by a particular response and is used to compute the SSI as

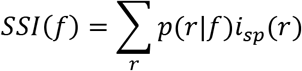

where *p*(*r*|*f*) is, for a particular frequency *f*, the fraction of responses equal to binned value *r*. The SSI is the average information present in the evoked responses when vibration frequency *f* is present. Significant SSI values therefore identify vibration frequencies well-encoded by the neuron.

The undersampled probability distributions, due to a finite number of stimulus repetitions at each frequency, result in biased SSI values. To correct for the bias, we computed the SSI with shuffled frequency labels and subtracted it from the SSI calculated with non-shuffled data. This procedure was repeated 1999 times and if the lower 99^th^ percentile of the bias corrected SSI values was not more extreme than zero, SSI was deemed significant (i.e. randomization test at significance level p<0.01).

### Tuning curve fitting

To characterize a neuron’s tuning to vibration frequency, the normalized mean responses were fit to frequency (*f*) by the following descriptive function using the method of non-linear least squares:

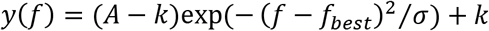

where σ = σ_ and *k* = 0 if *f* < *f*_best_, and σ = σ_+_ and *k* = *k*_+_ if *f* > f_best_. The five free parameters *A, k*_+_, σ_, σ_+_ and *f_best_* were fit by constraining *k_+_to* be less than 0.7 (normalized Δf/f_0_ to maximum mean response), *A* greater than 0.8 and 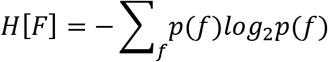 less than 3000 Hz using the non-linear least squares method. The fits were classified as: (i) frequency selective if *f_best_* ≤ 730 Hz and σ_+_ was not fixed at the bound, (ii) increases-to-plateau if *f_best_* ≤ 730 Hz and σ_+_ was fixed at the bound, in which case the fit was repeated with unbounded σ_+_, and (iii) monotonically increasing if *f_best_* > 730 Hz.

### Phase-locked spiking

To determine whether a neuron was entrained by the sinusoidal vibration, we first calculated for each spike its corresponding phase in the stimulus cycle (Fig. 4B) and grouped phase values across stimulus repetitions. For a given stimulus frequency, spiking was deemed to be phase-locked if the null hypothesis that phase angles are uniformly distributed could be rejected at significance level 0.01 (Rayleigh test for non-uniformity of circular data). Second, to control for variability in stimulus/response onset times, we calculated standardized inter-spike-intervals. For a given vibration frequency, inter-spike-intervals (*ISI*) of all possible spike pairs that occurred during stimulation were calculated and grouped across stimulus repetitions. Because entrained spiking should yield an *ISI* distribution that peaks at integer multiples of the sinusoidal stimulus period *T*, values were converted to standardized inter-spike-intervals (*S_ISI_*) according to:

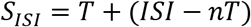

where *nT* is the integer multiple of *T* closest to *ISI*. Spiking was deemed to be entrained by the stimulus periodicity if the null hypothesis that *S_ISI_* values are uniformly distributed could be rejected at significance level 0.01 (χ^2^ test).

## Video analysis

For optogenetic and touch stimuli, videos of the mouse forepaw were continuously acquired with a 1/3” CMOS camera (Firefly MV FMVU-03MTM, Point Grey Research). A custom video acquisition software programmed in Matlab (Mathworks) saved 8-bit greyscale images (376 × 240 pixels) to disk at a rate of 30 frames/s. The start of trial TTL pulses issued by the behavior PC were received as external triggers and frame times relative to trigger onset were saved in the image headers. The saved timestamps were used for temporal alignment with Ca^2+^ traces during post-hoc analysis.

Images corresponding to frames immediately preceding stimulus onset were used for estimating the stimulated paw region. The horizontal and vertical means of each image were concatenated and the obtained traces mapped into a two-dimensional space with t-Distributed Stochastic Neighborhood Embedding (t-SNE) (van der Maaten and Hinton, 2008b) using the t-SNE Matlab toolbox (lvdmaaten.github.io/tsne). The initial dimensionality reduction via Principal Component Analysis performed by t-SNE was set to 5 dimensions and the t-SNE perplexity parameter to 15. The mapped points formed identifiable clusters, each corresponding to a different stimulated location on the paw (Fig. 7B). Paw locations associated with each cluster were subjectively determined by visual inspection. We estimated a neuron’s receptive field by color coding the mapped points based on evoked Δf/f_0_ responses (Fig. 7A-C and Fig. S7A-C).

## Statistics and data

No statistical methods were used to predetermine sample size and all experimental animals were included in the analysis. The normality assumption was tested with the Kolmogorov-Smirnov test. Non-parametric tests were used when the normality assumption was not met. All data analyses were performed with custom written routines in Matlab.

All data analysed and all custom Matlab code used for the analysis in the current study are available from the corresponding author upon request.

## Acknowledgments

We thank Vincent Hayward and Sliman Bensmaia for their advice and comments on the manuscript, Robert Zimmerman and Gregorio Galinanes for help with histology, Sylvain Crochet for advice on the electrophysiology experiments and Claudia Bonardi for help with training mice on the forelimb reaching task. This work was supported by the Swiss National Science Foundation (PP00P3_133710), the European Research Council (OPTOMOT), the New York Stem Cell Foundation and the International Foundation for Paraplegia Research. D.H. is a New York Stem Cell Foundation-Robertson Investigator.

---

**Figure s1.**
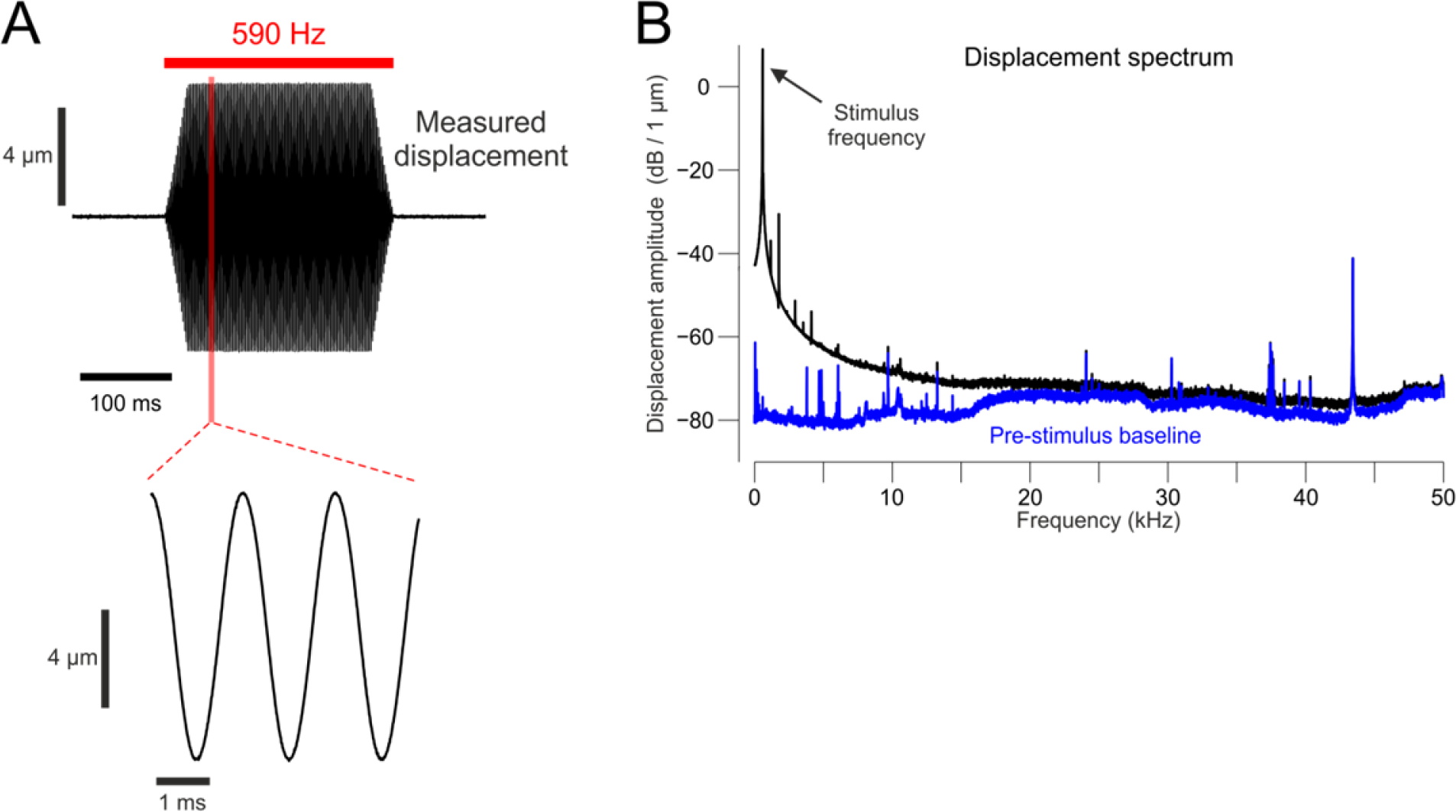
Spectral analysis of measured displacement produced by the vibrotactile stimuli. **(A)** Example of a displacement measured by the stimulator’s strain gauge (black) of a single 590 Hz vibration. The bottom trace shows an expansion in time of the interval highlighted in red. **(B)** Amplitude spectrum of the displacement produced by the 590 Hz stimulus indicates that the physical vibration consisted of a single frequency component (i.e. was a pure sinusoid). Components apparent at higher frequencies were highly attenuated relative to the stimulation frequency and were for the most part continuously present as revealed by the signal spectrum of the pre-stimulus baseline period (blue). Analogous amplitude spectra were obtained for stimuli at all other tested frequencies and amplitudes.

**Figure s2.**
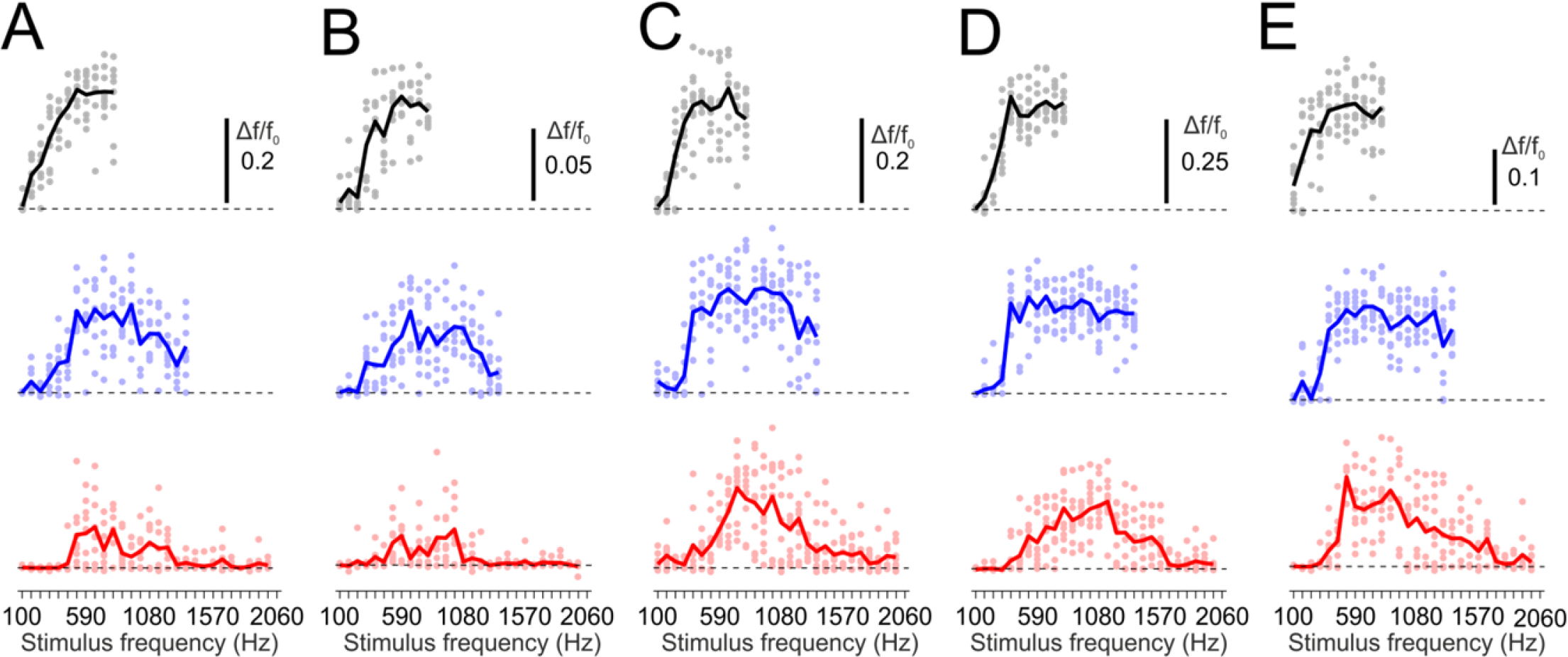
Tuning curves change with vibration amplitude. **(A-E)** Tuning curves changes and extension of the tested frequency range of five example neurons as the vibration amplitude is attenuated from 5.45 μm (black) to 1.08 μm (blue), to 0.23 μm (red).

**Figure s3.**
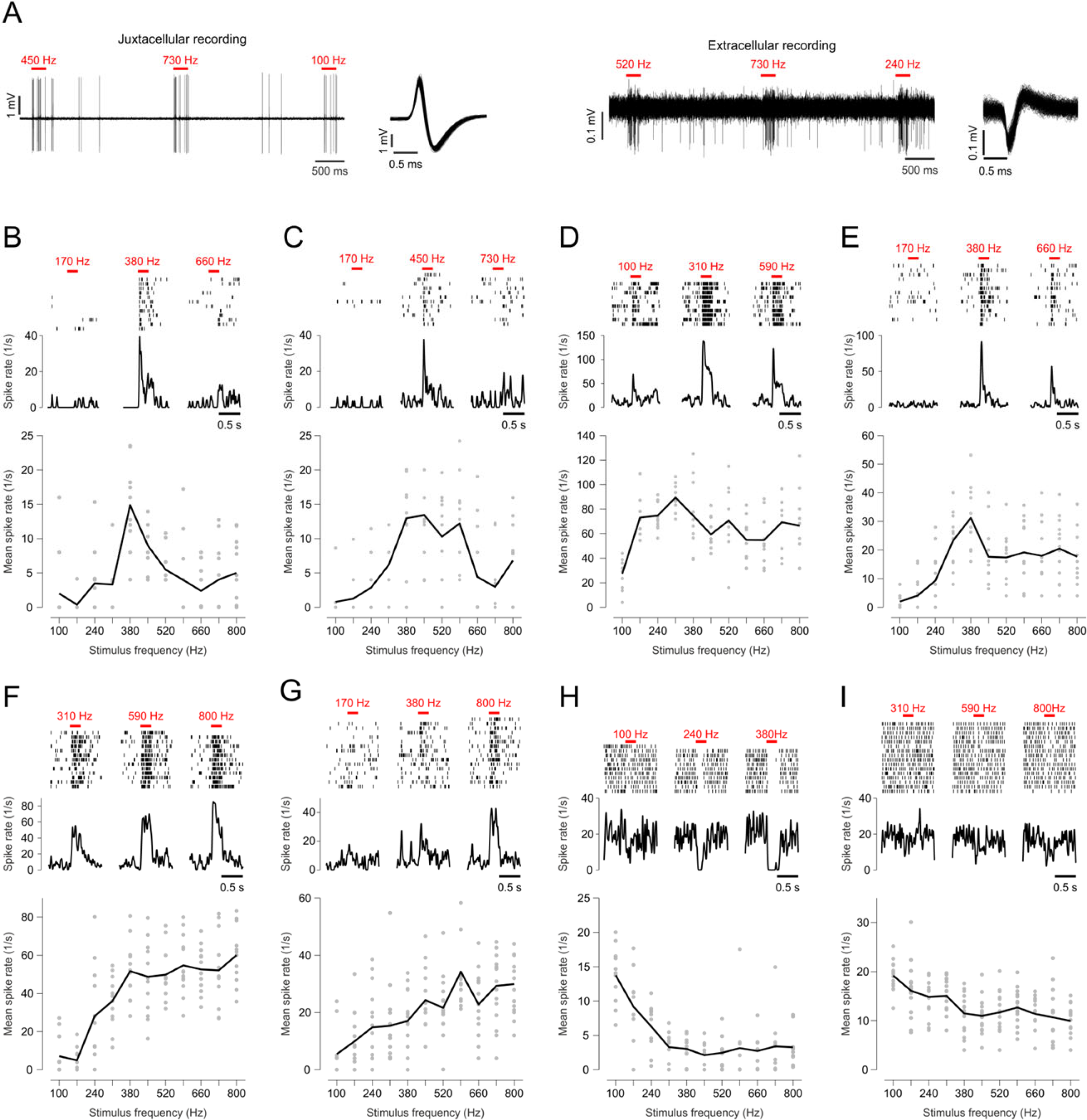
Electrode recordings of vibrotactile responses in fS1. **(A)** Examples of activity traces and action potentials recorded juxtacellularly and extracellularly in response to forepaw vibrations. **(B-I)** Spike rasters (top) and spike rate estimates (middle) of representative units recorded in fS1 in response to three chosen stimulus frequencies with their complete tuning curves based on the evoked mean spike rates (bottom).

**Figure s4.**
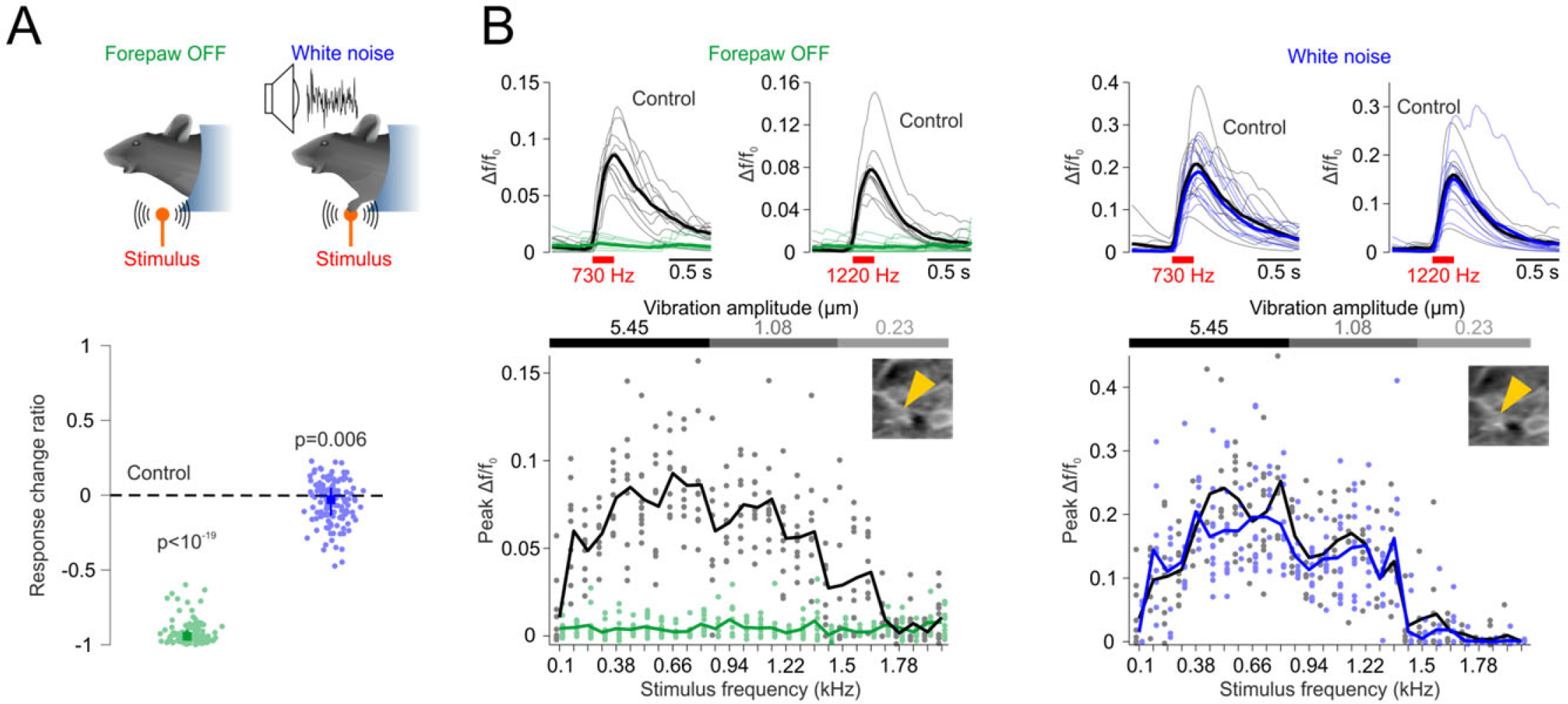
Absence of auditory responses in imaged neurons. **(A)** To test for the possibility that airborne sounds activated the imaged neurons in fS1, the forepaw was moved off the stimulator. To test for the possibility that vibrations propagated through the body to the cochlea, loud white noise was used to mask this hypothetical stimulation. The response change ratios (median ± quartiles) relative to the control condition (zero level) averaged across all frequencies with significant activity increases in the control condition show that with the forepaw off the stimulator, sound alone could not drive increases in Δf/f_0_ (p<10^−21^, Wilcoxon signed-rank test, n=113 cells, 9 mice) and that the white noise mask had but a small aversive effect on the responses (p=0.006, n=128 cells, 9 mice). Significant Δf/f_0_ increases in none of the individual neurons could be evoked by sound alone at any frequency (p<0.01, randomization test, n=1999 repetitions). **(B)** A representative fS1 neuron activated by forepaw vibrations (black: control condition) was irresponsive to sound alone (green: forepaw off condition) and unaffected by the auditory mask (blue: white noise condition). Top: denoised Δf/f_0_ responses evoked by 730 Hz and 1220 Hz vibrations (thin lines: individual stimulus responses, thick lines: mean response). Bottom: frequency tuning curves tested with four different vibration amplitudes, as depicted by the greyscale bar, to cover the extended range of frequencies (dots: individual responses, lines: mean response). Inset: cropped two-photon image depicting the same responding neuron in both conditions (yellow arrow).

**Figure s5.**
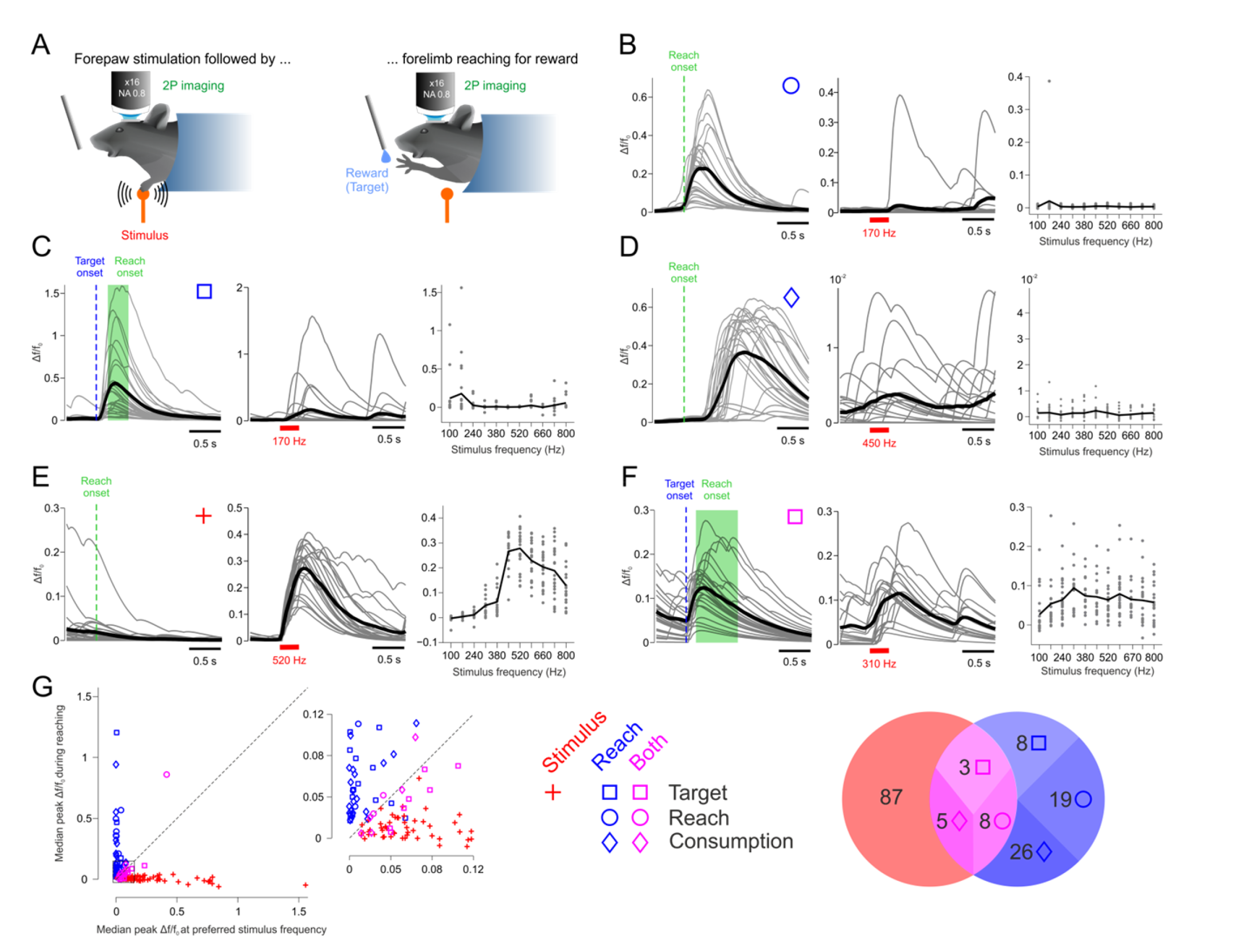
Forelimb movements do not account for the frequency tuned vibrotactile responses. **(A)** To test whether stimulus-locked motor events could drive neural responses in fS1, we trained mice to reach with their forelimb for the water reward following the series of forepaw vibrations on each trial. **(B-D)** Representative neurons responding to either reach onset (B), target onset (C) or around the time of water consumption (d) but not to forepaw vibrations. Left and middle: denoised Δf/f_0_ responses to reaching and a chosen vibration frequency, respectively. (thin lines: individual stimulus responses, thick lines: mean response, green shading in (C): range of reach onset times). Right: frequency tuning curves (dots: individual responses, lines: mean response). **(E)** Representative neuron responding to forepaw vibrations but not forelimb movements. **(F)** Representative neuron responding to both events but not showing frequency tuned activity. **(G)** Left: median peak Δf/f_0_ during reaching vs. at preferred vibration frequency for all imaged neurons (156 cells, 3 mice, 8 FOVs). Inset: enlarged view of the data in the black square. Right: depiction of numbers of neurons responding to vibrotactile stimulation (red cross), reaching onset, target onset or water consumption (blue symbols) or to both events (magenta symbols).

**Figure s6.**
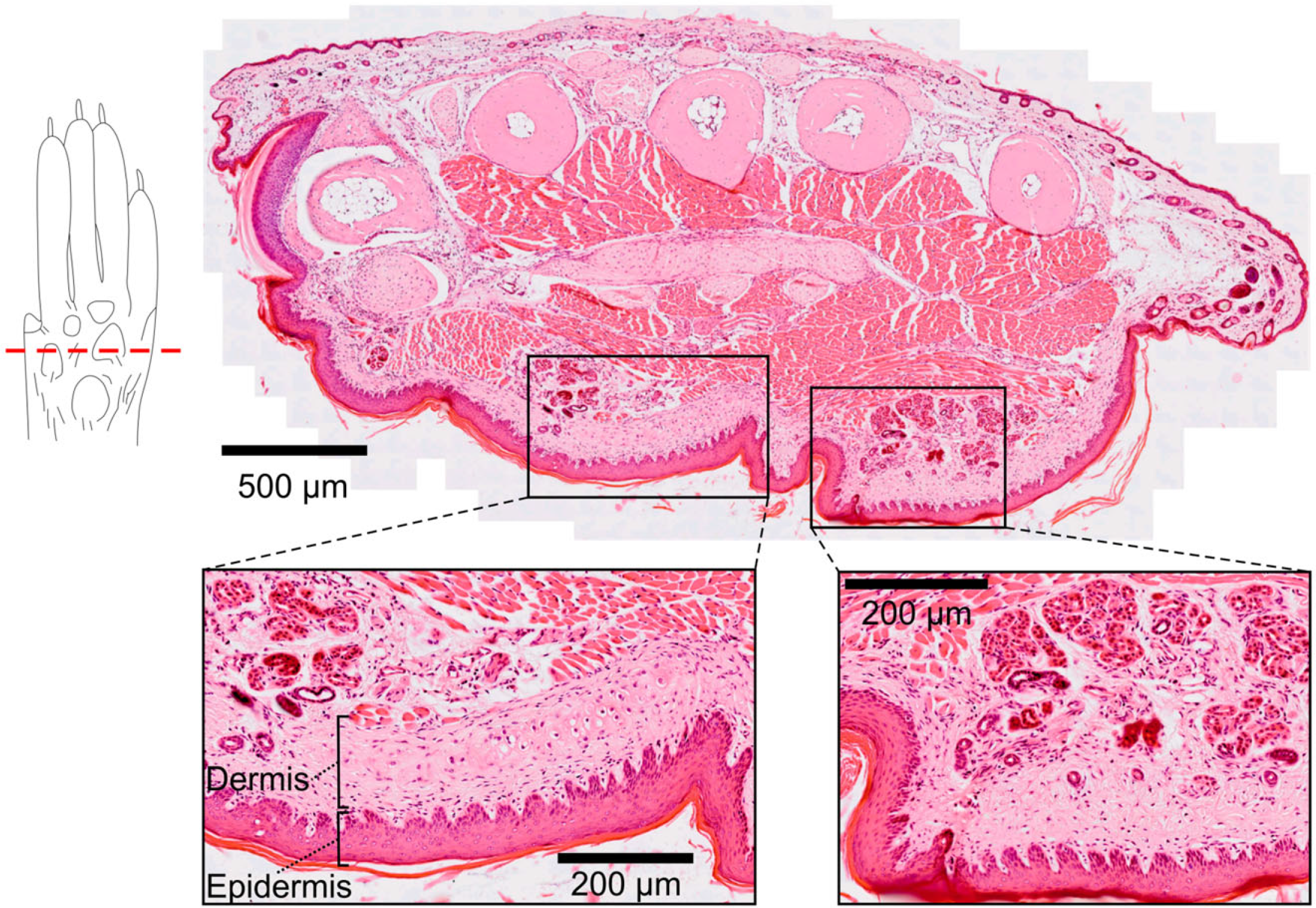
Absence of Pacinian corpuscles in the dermal layer of forepaw pads. A transverse 5 μm section (haematoxylin and eosin stained) of a mouse forepaw. The magnified panels show the absence of Pacinian corpuscles in the dermal layer of the palm pads.

**Figure s7.**
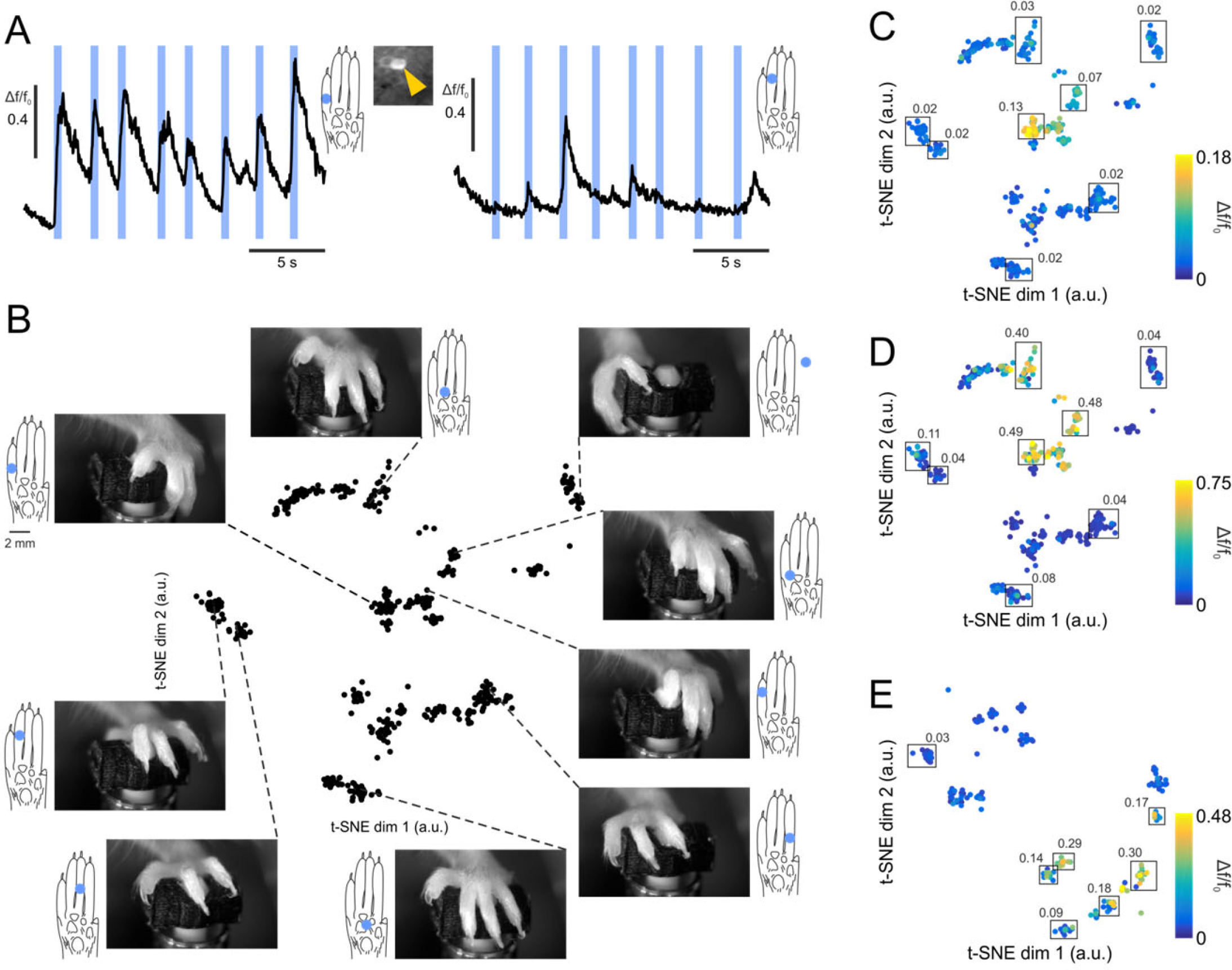
Optogenetic forepaw stimulation analysis. **(A)** Δf/f_0_ traces of an example neuron on two successive optogenetic stimulation (blue lines) trials with different paw placements relative to the optic fiber (blue filled circles). **(B)** Concatenated horizontal and vertical means of forepaw images immediately prior to stimulus onset were projected into a low-dimensional space using t-SNE. The formed clusters correspond to stimuli with highly similar paw placements relative to the optic fiber and were used to subjectively estimate locations of the optogenetic stimulation on the paw, as illustrated. **(C-E)** Δf/f_0_ responses for each neuron were used to color code the t-SNE projection points. The clearly visible clustering of high vs. low Δf/f_0_ values implies that stimulus locations on the paw are separable based on the size of Δf/f_0_ responses they evoke. The black squares denote clusters in t-SNE space used to evaluate average responses (indicated numerical values) evoked by stimulus locations shown in Fig. 6 (A-C).

**Movie S1. Vibratory stimulation of 1 mouse forelimb during cortical imaging**. Successive vibrations are delivered to the forelimb of a mouse followed by water reward. The numerical values indicate the moment of stimulation and the vibration frequency.

**Movie S2. Vibratory stimulation of mouse forelimb followed by goal-directed reaching**. Successive vibrations are delivered to the forelimb of a mouse followed by reaching for a water droplet. The numerical values indicate the moment of stimulation and the vibration frequency.

Movie S3. Optogenetic stimulation of mouse forepaw during cortical imaging.

Movie S4. Monitoring of paw placement during optogenetic stimulation.

